# The origin and evolution of *loqs2*: a gene encoding an antiviral dsRNA binding protein in *Aedes* mosquitoes

**DOI:** 10.1101/2021.12.21.473503

**Authors:** Carlos F. Estevez-Castro, Murillo F. Rodrigues, Antinéa Babarit, Flávia Viana Ferreira, Eric Marois, Rodrigo Cogni, João Trindade Marques, Roenick Proveti Olmo

**Affiliations:** Department of Biochemistry and Immunology, Instituto de Ciências Biológicas, Universidade Federal de Minas Gerais, Belo Horizonte, Brazil; Institute of Ecology and Evolution, University of Oregon, Eugene, Oregon, United States; CNRS UPR9022, Inserm U1257, Université de Strasbourg, Strasbourg, France; Department of Ecology, Institute of Biosciences, University of São Paulo, São Paulo, Brazil

**Author notes:** **Correspondence:** Roenick Proveti Olmo -, João Trindade Marques.

**Keywords:** *loqs2*, RNA interference, *Aedes* mosquitoes, double-stranded RNA binding proteins (dsRBP)

## Abstract

Mosquito borne viruses such as dengue, Zika, yellow fever and Chikungunya cause millions of infections every year. These viruses are mostly transmitted by two urban-adapted mosquito species, *Aedes aegypti* and *Aedes albopictus*, that appear to be more permissive to arbovirus infections compared to closely related species, although mechanistic understanding remains unknown. *Aedes* mosquitoes may have evolved specialized antiviral mechanisms that potentially contribute to the low impact of viral infection. Recently, we reported the identification of an *Aedes* specific double-stranded RNA binding protein (dsRBP), named Loqs2, that is involved in the control of infection by dengue and Zika viruses in *Ae. aegypti*. Loqs2 interacts with two important co-factors of the RNA interference (RNAi) pathway, Loquacious (Loqs) and R2D2, and seems to be a strong regulator of the antiviral defense. However, the origin and evolution of *loqs2* remains unclear. Here, we describe that *loqs2* likely originated from two independent duplications of the first dsRNA binding domain (dsRBD) of *loquacious* that occurred before the radiation of the *Aedes Stegomyia* subgenus. After its origin, our analyses suggest that *loqs2* evolved by relaxed positive selection towards neofunctionalization. In fact, *loqs2* is evolving at a faster pace compared to other RNAi components such as *loquacious, r2d2* and *Dicer-2* in *Aedes* mosquitoes. Unlike *loquacious*, transcriptomic analysis showed that *loqs2* expression is tightly regulated, almost restricted to reproductive tissues in *Ae. aegypti* and *Ae. albopictus*. Transgenic mosquitoes engineered to ubiquitously express *loqs2* show massive dysregulation of stress response genes and undergo developmental arrest at larval stages. Overall, our results uncover the possible origin and neofunctionalization of a novel antiviral gene, *loqs2*, in *Aedes* mosquitoes that ultimately may contribute to their effectiveness as vectors for arboviruses.

## Introduction

*Aedes aegypti* and *Aedes albopictus* are the major vectors for arthropod-borne viruses (arboviruses), such as yellow fever, dengue (DENV), Zika (ZIKV), and chikungunya (CHIKV) viruses. DENV alone is estimated to be responsible for approximately 400 million infections and 20,000 deaths per year worldwide [1]. Globalization, urbanization and climate change are contributing to the spread of *Aedes* mosquitoes to previously uncolonized regions impacting virus transmission and emergence of new arboviruses with potential to affect human health [2]. Despite the urgent need, there are no treatments or vaccines for most arboviral diseases [3].

*Ae. aegypti* and *Ae. albopictus* are not only well adapted to urban settings and anthropophilic feeding but also seem to be naturally more susceptible to arboviruses under laboratory conditions [4–7]. This suggests that these mosquito species may be intrinsically more susceptible to arboviruses, which is a big part of their overall vector competence [7]. It remains unclear whether there are clear genetic markers of vector competence in *Aedes* mosquitoes, especially compared to other mosquito species such as *Culex* and *Anopheles*. One possibility is that the antiviral defense in *Aedes* mosquitoes diverged from closely related species, allowing virus infection without significant harm to the insect host [8–10].

RNA interference (RNAi) and, in particular, the small interfering RNA (siRNA) pathway, is a broad antiviral defense mechanism in insects [9,11,12]. Virus-derived double-stranded RNAs (dsRNA) are processed by Dicer-2 (Dcr-2) into siRNAs that are subsequently loaded into the nuclease Argonaute-2 (AGO2) to form the RNA-induced silencing complex (RISC). This complex is then able to target complementary viral RNAs and thus inhibit virus replication. RNAi co-factors such as the dsRNA binding proteins (dsRBPs) Loqs and R2D2 are also essential for the biogenesis and loading of small RNAs to the RISC complex [13] and, in the *Ae. aegypti* cell line Aag2, Loqs and R2D2 also seem to act non-redundantly in the antiviral branch of the siRNA pathway [14]. The arms race between viruses and antiviral RNAi resulted in rapid evolutionary rates of siRNA pathway genes such as *Dcr-2, AGO2* and *r2d2*. This adaptive signature has been observed in *Drosophila* and mosquito species [15–17]. In addition, duplications and losses of RNAi genes are observed in multiple insect lineages, resulting in great RNAi divergence and specialization even between related species [18]. Indeed, we have recently identified a novel dsRBP gene in *Aedes* mosquitoes, *loqs2*, that is likely a paralog of *loqs* [19]. Silencing of *loqs2* showed significantly increased levels of DENV RNA during systemic infection and, in addition, ectopic expression of *loqs2* in the midgut resulted in the control of DENV and ZIKV infection in the mosquito [19].

Here, we used different bioinformatic approaches to unveil the origin and evolution of *loqs2*. Our results indicate that *loqs2* is a result of independent duplications of the first dsRNA binding domain of *loqs* that occured before the *Aedes Stegomyia* subgenus radiation. *loqs2* underwent relaxed positive selection after its origin and is evolving at a faster pace than other RNAi components such as *loqs* and *Dicer-2* in *Aedes* mosquitoes. We observed that *loqs2* expression is tightly regulated in *Ae. aegypti* and *Ae. albopictus* and that ectopic ubiquitous expression caused developmental arrest at the larval stages. Taken together, our results corroborate the hypothesis that *loqs2* has evolved through neofunctionalization within some *Aedes* species to regulate different aspects of the antiviral response.

## Results

### *loqs2* is a paralog of *loqs* that originated in the ancestor of the *Aedes Stegomyia* subgenus

To analyze the origin of *loqs2*, we screened the genomes of flies and mosquitoes for proteins with at least one double-stranded RNA-binding domain (dsRBD) (InterPRO accession IPR014720). These were evaluated to determine their phylogenetic relationships (**Figure 1A**, extended version in **Supplementary Figure 1A**). The overall topology of the tree of dsRBPs shows monophyletic origins for each protein ortholog identified, with Loqs2 proteins located as a single sister clade of Loqs. These results confirm that *loqs2* is a paralog of *loquacious* (*loqs*), a component of the RNAi pathway.

**Figure 1.**
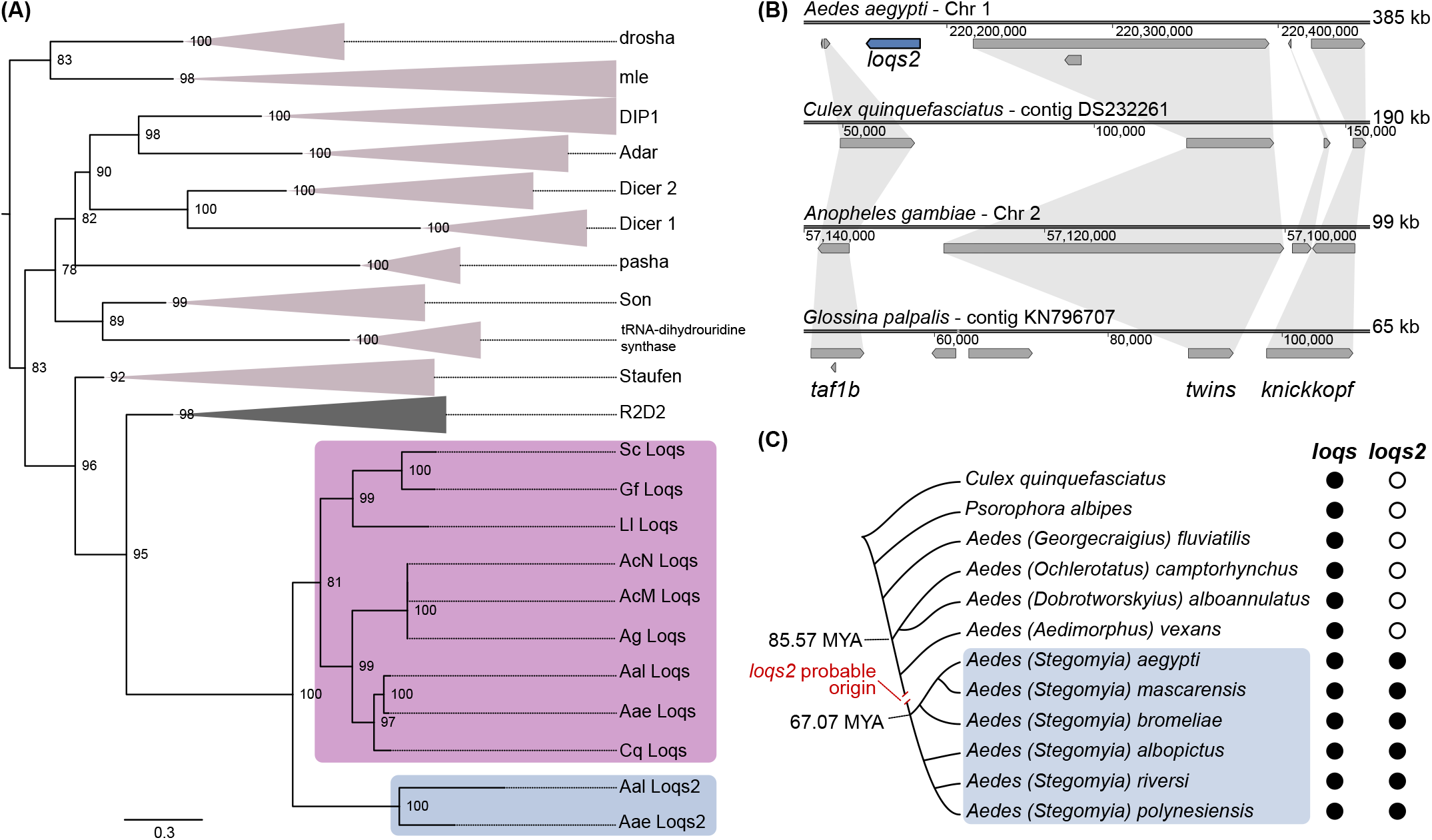
*loqs2* is a dsRBP paralog of *loqs* that originated in the ancestor of the *Aedes Stegomyia* subgenus. **(A)** Maximum likelihood phylogenetic tree constructed with the amino acid sequences of the dsRNA-binding proteins (dsRBPs) found in the genomes of *Ae. aegypti* (Aae), *Ae. albopictus* (Aal), *Culex quinquefasciatus (*Cq*), Anopheles gambiae* (Ag), *Anopheles coluzzii* Mali-NIH (AcM), *Anopheles coluzzii* Ngousso (AcN), *Lutzomyia longipalpis* Jacobina (Ll), *Stomoxys calcitrans* USDA (Sc) and *Glossina fuscipes* IAEA (Gf). Branches corresponding to the orthologs of drosha, mle, DIP1, Adar, Dicer-2, Dicer-1, pasha, Son, tRNA-dihydrouridine synthase, Staufen and R2D2 are collapsed to facilitate visualization. Tree is rooted at the midpoint for visualization purposes. Node values correspond to 1000 ultra-fast bootstrap iterations. A non-collapsed tree is available at **Supplementary Figure 1A. (B)** Synteny analysis of *loqs2* flanking genomic regions among *Ae. aegypti, C. quinquefasciatus, An. gambiae* and *G. palpalis*. **(C)** Schematic cladogram indicating the probable origin of *loqs2*. The putative appearance of *loqs2* was inferred from the identification of sequences matching *loqs2* coding sequence in high-throughput sequencing data publicly available and are illustrated in **Supplementary Figure 1B**. Phylogenetic relationships and molecular clock were extracted from Wilkerson et al. [23] and da Silva et al. [22].

Interestingly, we also identified independent cases of gene duplications among the other species and ortholog groups, reinforcing the importance of dsRBPs. The dsRBP divergence and specialization among the species evaluated here also suggest that dsRNA recognition has been tuned by the individual evolutionary history towards specialization, even among closely related species. The clear monophyly found among ortholog groups identified, also shown by the relationship between the Loqs and Loqs2 clade, showed that despite fast diversification there is a functional convergence potentially to maintain key properties. An example is the traceability of dsRBPs from apparently simple organisms to very complex ones, such as the orthologs *TRBPs/PACT* in vertebrates and urochordates, *r2d2*/*loqs* in arthropods, *RDE-4* in nematodes, and *DRB4* in plants, or also *Dicer* ortholog genes in all aforementioned clades [20]. Of note, it is compelling to expect more duplication events among genes containing dsRBDs that are involved in the antiviral arm of RNAi, as they are expected to be constantly under evolutionary pressure due to their role in immunity [16]. However, we also observed paralog duplications among dsRBD-containing genes that perform housekeeping functions such as the microRNA pathway, which is well recognized to be evolving under strong purifying selection [15,16].

Considering that *loqs2* is only found in *Aedes* mosquitoes, we evaluated the corresponding syntenic regions surrounding *loqs2* in other dipteran species using available genomes at Vectorbase (**Figure 1B**). This analysis showed genetic conservation of synteny structure in the region, in which three genes (*taf1b, twins*, and *knickkopf*) were present in all dipteran species we analyzed, from *Ae. aegypti* to *Glossina palpalis*. Interestingly, other than the presence of *loqs2* in *Ae. aegypti*, we observed other differences in this genomic region between dipteran species. Two genes appeared between *taf1b* and *twins* in *G. palpalis* that are absent in the other dipterans studied. In addition, *Anopheles gambiae, Culex quinquefasciatus* and *Ae. aegypti* possess an extra conserved gene inserted between *twins* and *knickkopf*, which is absent in *G. palpalis*. This suggests that this region might be prone to gene insertions and deletions. Our analysis also highlights the large genomic expansion that *Ae. aegypti* has undergone (**Figure 1B**) [21]. The syntenic genomic region from the left and rightmost genes in *Ae. aegypti* is twice as large compared to *C. quinquefasciatus* and five times larger compared to *G. palpalis*.

To further trace the origin of *loqs2* in the *Aedes* genus, we analyzed publicly available high-throughput RNA-seq libraries and whole genome sequencing (WGS) data from various species. Data included four species belonging to different subgenus (*i*.*e*., *Georgecraigius, Ochlerotatus, Dobrotworskyius, and Aedimorphus*), and four species from the *Stegomyia* subgenus. We also analyzed RNA-seq data from the mosquito *Psorophora albipes*, species closely related to the *Aedes* genus (**Figure 1C**). Here we used reference genes from *Ae. aegypti* and *Ae. albopictus* to map libraries of the other mosquitoes, to identify potential orthologous genes. As a control, we were able to detect sequences mapping to the *loqs* gene from *Ae. aegypti* or *Ae. albopictus* in all nine datasets (**Supplementary Figure 1B**). In contrast, we only observed sequences mapping to the *Ae. aegypti* and *Ae. albopictus loqs2* genes in libraries derived from species in the *Stegomyia* subgenus. Furthermore, sequence coverages were smaller than that observed for *loqs*, which is expected considering the interspecific diversification of *loqs2*. Altogether, our results suggest that *loqs2* originated by a duplication of the *loqs* gene that occurred before or at the radiation of the *Aedes Stegomyia* subgenus, which is estimated to have occurred between 85.57 to 67.07 million years ago (MYA) (**Figure 1C**) [22]. Such estimation ranges from the late Cretaceous, passing the K-PG mass extinction, up to the beginning of the Paleocene. Thus, the *Aedes* radiation that followed the K-PG mass extinction might have also influenced the origin and evolution of *loqs2* among the *Stegomyia* subgenus. Apart from *Ochlerotatus* and *Dobrotworskyius*, we did not find *loqs2* in the *Aedimorphus* subgenus. This subgenus is phylogenetically closer to the *Stegomyia* than the other two [23]. Thus, we suspect *loqs2* originated near to the last common ancestor of the *Stegomyia* (67.07 MYA). However, as there is no available dating time of the separation between *Aedimorphus* and *Stegomyia* we cannot further narrow down the estimated origin of *loqs2*.

### *loqs2* originated from two independent *loqs* dsRBD1 duplication events

Similar to *loqs* and *r2d2*, the *loqs2* coding sequence is expected to produce a small protein containing two canonical dsRBDs (**Figure 2A**). However, the Loqs2 protein differs from Loqs by the distance between the two dsRBDs and the absence of a third domain, called Staufen C (ST), which is a dsRBD-like predicted to bind to other proteins [24]. Such differences in domain organization prompted us to investigate more carefully the origins of *loqs2* by interrogating its structural features. First, we explored the phylogenetic relationships between individual dsRBDs from *loqs, loqs2* and *r2d2*. Although our previous data showed *loqs2* is a paralog of *loqs*, we also included *r2d2* domains to rule out any possible relationship between those proteins. We observed that both the first and second dsRBDs of *loqs2* (D1, D2) grouped together with the first dsRBD of *loqs* but relatively far from the second dsRBD. Local alignment of the dsRBDs and Staufen C (ST) amino acid sequences of Loqs and Loqs2 from *Ae. aegypti* and *Ae. albopictus* (**Figure 2B**) confirmed that both Loqs2 dsRBDs (Loqs2-D1 and Loqs2-D2) were more similar to the first dsRBD of Loqs (Loqs-D1). Importantly, this analysis also showed that they are more similar to Loqs-D1 than to each other (**Figure 2B**) and highlighted the great conservation seen among the evaluated *loqs* orthologs compared to the interspecific diversification of *loqs2* genes (**Supplementary Figure 1B**). We also compared structural properties of Loqs and Loqs2 dsRBDs by using the resolved three-dimensional structures of the dsRBD1 and dsRBD2 from Loqs-PD of *D. melanogaster* (PDB ids 5NPG and 5NPA, respectively) as a reference [25]. We first predicted *in silico* the structures of individual dsRBDs of Loqs and Loqs2 from *Ae. aegypti* and *Ae. albopictus* and used the best model for comparisons using the DALI server [26]. Both dsRBDs from Loqs2 were more structurally related to the first dsRBD of Loqs (**Figure 2C**). Notably, the second dsRBD of Loqs presented low structural similarity to all the other domains analyzed (Loqs-D1, Loqs2-D1 and Loqs2-D2). Taken together, our findings suggest that *loqs2* originated from two independent tandem duplications of the first dsRBD of *loqs* and not a direct gene duplication. By using *Ae. aegypti* as a model for genome organization in the *Stegomyia* subgenus, we can suggest that such domain duplications from the *loqs* gene located in chromosome 2 were then inserted into chromosome 1, where *loqs2* is now present (**Figure 2D**).

**Figure 2.**
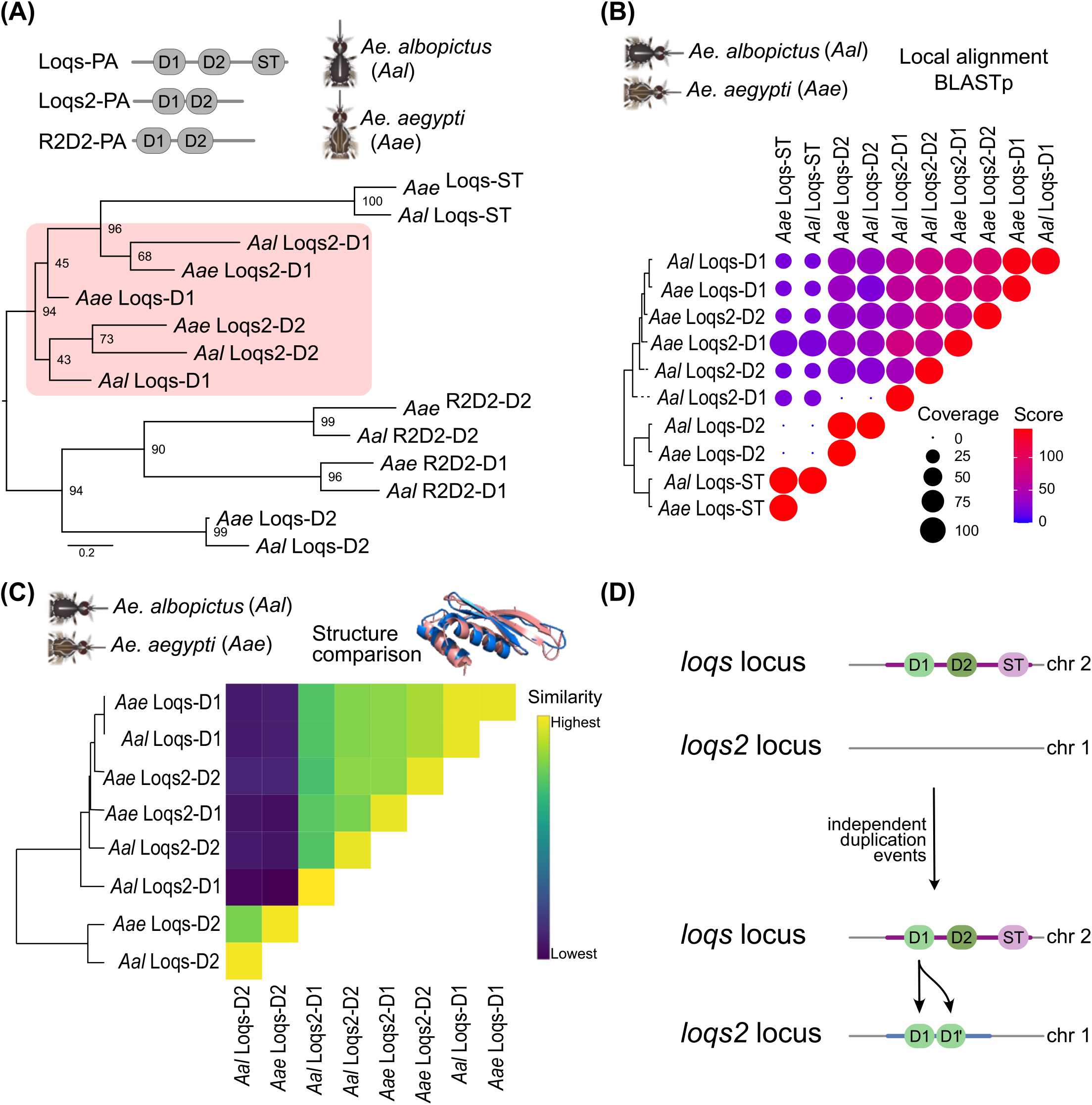
*loqs2* originated from two independent *loqs* dsRBD1 duplication events. **(A)** Phylogenetic relationships among the *Ae. aegypti* and *Ae. albopictus* Loqs2, Loqs, and R2D2 double-stranded RNA-binding domains (dsRBDs) D1 and D2, and Staufen C (ST) domain. Phylogeny was inferred by maximum likelihood and tree is rooted at the midpoint for visualization purposes. Node values correspond to the result of 1000 ultra-fast bootstrap iterations. **(B)** Correlation matrix showing BLASTp alignment score and coverage results from the comparison of *Ae. aegypti* Loqs and Loqs2 dsRBD amino acid sequences. **(C)** Heatmap showing the structural comparison between Loqs and Loqs2 dsRBDs D1 and D2 models using DALI server. **(D)** putative model for *loqs2* origin.

### Relaxed positive selection shaped *loqs2* evolution towards neofunctionalization

Gene duplications can have a range of outcomes. Purifying selective constraints are expected to pressure one copy to maintain its original functions, while the other is free to accumulate mutations. In this scenario, the most likely outcome is pseudogenization of one copy. Alternatively, accumulation of mutations can lead to subdivision of ancestral functions between duplicated genes, called subfunctionalization. Occasionally, mutations may lead to neofunctionalization that, if advantageous, will drive fixation of the new gene in the population [27]. We hypothesized that the appearance of *loqs2* might have had little impact on the function of *loqs* (**Figure 3A**), since key functions of Loqs are still conserved even in the presence of Loqs2 [14]. In addition, there is some evidence for the neofunctionalization of *loqs2* [19]. To evaluate the type of selections that shaped the evolution of *loqs* and *loqs2*, we estimated the ratio of nonsynonymous to synonymous substitutions (ω) using a codon-based phylogenetic framework. This allowed us to build a topology by maximum likelihood inference using dipteran species, evaluating *loqs* and *loqs2* as ingroups, and *r2d2* as outgroup (**Figure 3B**). First, we performed a test for relaxation or intensification of selection using the software RELAX [28]. RELAX is based on the selection intensity parameter “*k”*, which establishes a comparison between the test and the reference branches of the phylogenetic tree. *k* > 1 indicates intensified selection while *k* < 1 indicates relaxed selection of test branches compared to the reference. We applied this strategy to *Aedes loqs* or *loqs2* clades as test branches compared to *loqs, loqs2* or *r2d2* clades as references. Our results showed that *loqs2* is evolving under relaxed selection compared to either *loqs* (*k* = 0.07) or *r2d2* (*k* = 0.09) (**Figure 3C**). In contrast, *loqs* is evolving under intensified selection compared to *loqs2* (*k* = 7.19) or *r2d2* (*k* = 2.82). These results support the hypothesis that *loqs* and *loqs2* are evolving under different selective regimes.

**Figure 3.**
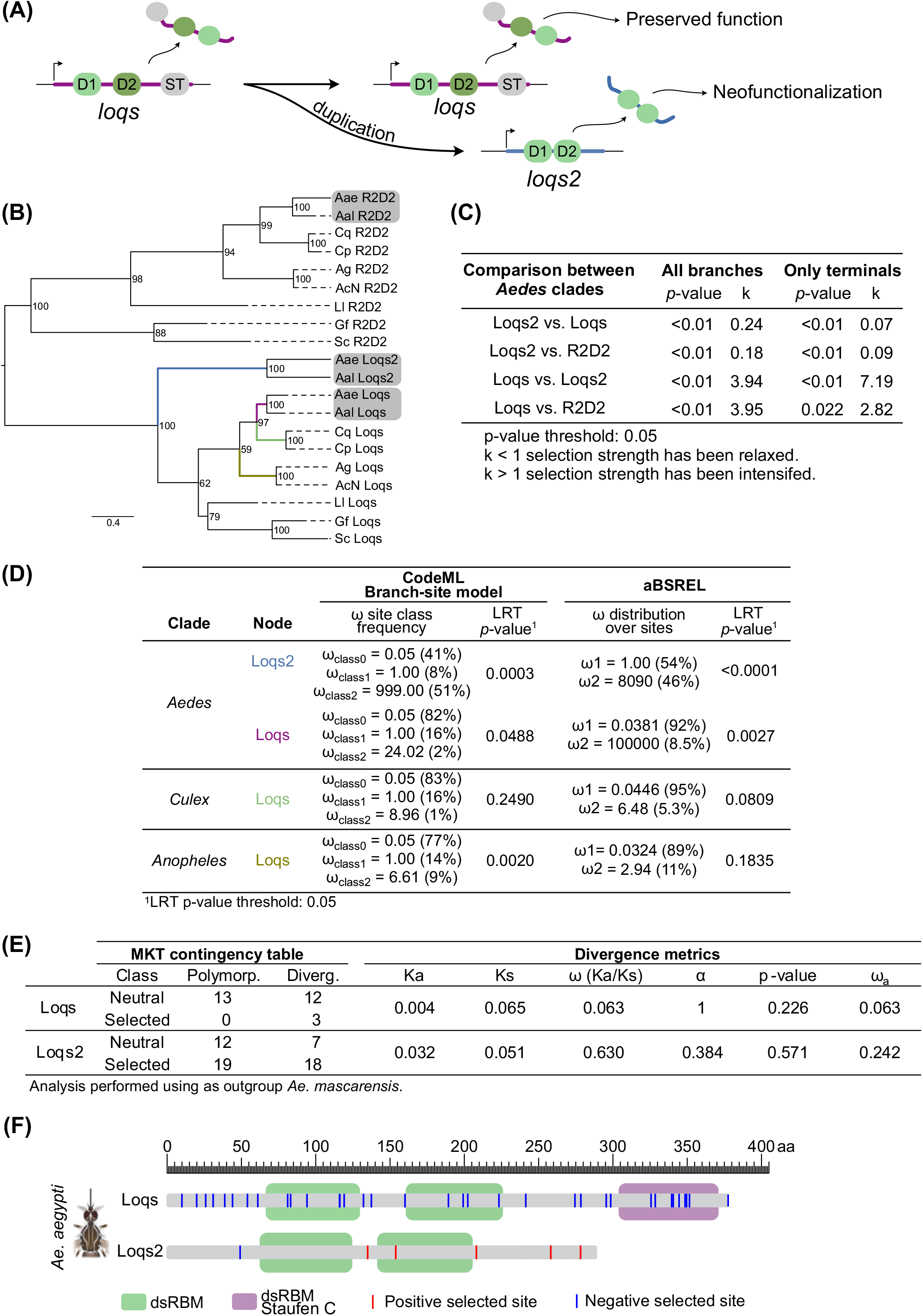
Relaxed positive selection shaped *loqs2* evolution towards neofunctionalization. **(A)** Proposed model of the evolutionary path followed by *loqs* and *loqs2* after the duplication events. **(B)** Maximum likelihood phylogenetic tree constructed with the coding sequences of *loqs, loqs2* and *r2d2* orthologs from *Ae. aegypti* (Aae), *Ae. albopictus* (Aal), *C. quinquefasciatus* (Cq), *C. pipiens* (Cp), *An. gambiae* (Ag), *An. coluzzii* Ngousso (AcN), *L. longipalpis* Jacobina (Ll), *S. calcitrans* USDA (Sc) and *G. fuscipes* IAEA (Gf). Tree is rooted at the midpoint for visualization purposes. Node values correspond to the result of 1000 ultra-fast bootstrap iterations. **(C)** Test for relaxation or intensification of selection performed using RELAX. **(D)** Test of selection using the branch-site-model implemented in CodeML and abSREL. **(E)** Divergence metrics and McDonald-Kreitman test (MKT) results from an *Ae. aegypti* population from a forest in Senegal compared to *Ae. mascarensis*. **(F)** Comparative scheme between *Ae. aegypti* Loqs and Loqs2 proteins. Sites represented in red or blue correspond to common amino acids under selection among *Ae. aegypti* and *Ae. albopictus*.

The unusual origin of *loqs2* may have favored an evolutionary scenario under relaxed selection that continues to the present date, despite *loqs2* origin dated of more than 67 MYA. We aimed here to predict the impact of *loqs2* appearance on selective pressures acting on *loqs*. However, to our understanding, this analysis undergoes conflicting aspects. On one hand, because of prominent differences between both proteins, one could expect that the appearance of *loqs2* might have had little impact on the original selective pressures acting on *loqs*. In this scenario, Loqs should conserve original key functions before and during the evolution of *Stegomyia* subgenus. On the other hand, previous results from our group show that Loqs2 interact with Loqs [19]. This suggests that the appearance of *loqs2* might have influenced *loqs*, affecting original selective pressures. Altogether, our data favor a hybrid hypothesis, where Loqs2 had little influence on Loqs evolution. However, to provide a stronger conclusion we would need to evaluate other *Aedes* species, whose genomes are not yet available while writing this manuscript. Thus, while *loqs* might have evolved under intense selection to maintain canonical functions [14], *loqs2* is evolving under relaxed selection, which allows diversification and, eventually, innovation **(Figure 3A)**.

The type of natural selection acting on a protein can be very useful to understand its functional evolution. For example, a gene evolving towards pseudogenization (loss of function) is expected to evolve under neutral selection, while a gene moving towards neofunctionalization is expected to present sites under positive/negative selection. To investigate the modes of selection acting on both genes, we used two complementary approaches: the branch-site-model implemented in CodeML (CodeML-BSM) [29], and abSREL [30]. Both methods allow the estimation of ω variations among sites and branches, but with different assumptions. Conceptually, the major difference between both methods is that CodeML permits only four specific configurations of branch-specific rate parameters [31], and abSREL infers the optimal number of rate categories for each foreground branch and allows any of the sites of the branch to evolve under any of these categories [30]. As we were mainly interested in the evolution of *loqs* and *loqs2* in the *Aedes* genus, instead of separately testing the terminals for each of the genes (*e*.*g*., Aae-*loqs2* or Aal-*loqs2)*, we tested branches before the species split, being the closest representatives of the evolutionary history of these proteins at the genus level. Interestingly, both methodologies found significant evidence of positive selection in *loqs* and *loqs2* branches in the *Aedes* genus (**Figure 3D**). Also, both methodologies found similar ω distribution over sites. In the case of the *Aedes loqs*, most sites showed signs of purifying selection (CodeML-BSM site class 0 = 82%, abSREL ω_1_= 92%), and a smaller frequency of sites show signs of positive (CodeML-BSM site class 2a-b= 2%, abSREL ω_2_=8.5%) or neutral selection (CodeML-BSM site class 1 = 16%). In contrast, roughly half of sites of *loqs2* showed evidence of positive selection (CodeML-BSM site class 2a-b= 51%, abSREL ω_2_=46%), while the other half had signs of purifying (CodeML-BSM site class 0 = 41%) or neutral selection (CodeML-BSM site class 1 = 8%, abSREL ω_1_= 54%). These results are consistent with the hypothesis of relaxed selection after the duplication that gave rise to *loqs2*, which was then followed by episodic adaptive evolution (**Figure 3C**).

To better understand the selective forces acting on *loqs* without the presence of *loqs2*, we also analyzed the *loqs* branches corresponding to *Culex* and *Anopheles* (**Figure 3D**). We did not find consistent evidence of positive selection considering both methods. Since the *Culicini* diverged around 130.49 MYA from the *Aedini*, and the *Anophelinae* diverged from both tribes around 182.75 MYA [22], a comparison of the *Aedes loqs* with the *Culex* ortholog is more reliable. Interestingly, this suggests that *loqs* experienced changes in its selective regime following *loqs2* origin and diversification. Possibly, the duplication event that gave rise to *loqs2* may have freed specific sites in *loqs* from previous constraints thus allowing positive directional selection to occur.

We also explored the recent evolutionary history of *loqs* and *loqs2* using the McDonald-Kreitman test (MKT) [32] available in the iMKT web server [33]. MKT infers the presence of recurrent positive selection by comparing polymorphic sites among a population to divergent sites between the population against an outgroup. Thus, the MKT examines the evolutionary period between the divergence of the outgroup species and the present state of the ingroup population. We used publicly available exome data from *Ae. aegypti* samples from a forest in Senegal [34] compared to *Aedes mascarensis* as an outgroup [35]. Our results showed that *loqs2* has more amino acid substitutions within the Senegal forest population (selected polymorphisms) or compared to *Aedes mascarensis* (selected divergences), than intra-or interspecific neutral substitutions. On the contrary, *loqs* mainly presented neutral substitutions with only a few divergent sites between both species. The rate of nonsynonymous divergence (*Ka*) to the proportion of neutral fixations *(Ks)* suggested that *Ae. aegypti loqs2 (Ka/Ks* = 0.63) is evolving substantially faster than *loqs (Ka/Ks* = 0.063) (**Figure 3E** and **Supplementary Figure 2A**). Although we observed an elevated *Ka/Ks* ratio for *loqs2*, especially when compared with *loqs*, it was not greater than 1, which would provide undeniable evidence of positive selection. In fact, the MKT results did not show significant evidence of positive selection neither for *loqs* nor *loqs2*. This lack of significance is likely due to the limitations of our exome dataset since we used a small mosquito population (n = 9) from Senegal as ingroup and a laboratory strain *Ae. aegypti* Liverpool as reference, which certainly does not reflect the diversity of current mosquito populations. Nonetheless, when we compared the *Ka/Ks* ratio of *loqs2* to the ratio of *dicer2*, a hallmark gene of positive selection in insects [16], including *Ae. aegypti* [17], we found that *loqs2* is also evolving considerably faster. In a similar fashion, fast evolving genes such as *D. melanogaster Dcr-2, AGO2* and *r2d2* display similar ω average values comparable to *loqs2* [36]. Of note, we have not compared the evolutionary rates of *Aedes loqs* to its *D. melanogaster* ortholog since it is not required for the antiviral siRNA response in the latter [37,38]. Altogether, our results indicate that *loqs2* is diversifying faster than *loqs* or *dicer-2* in a recent evolutionary scale. Furthermore, analysis of population genetics corroborates that *loqs* and *loqs2* are evolving under different selective regimes. In summary, both long-term and short-term evolutionary histories suggest that *loqs2* has evolved under relaxed positive selection and diversified towards neofunctionalization. Thus, *loqs2* could be fulfilling roles (*e*.*g*., antiviral immunity, control of gene expression) that *loqs* had not sufficiently evolved to sustain due to its shared role between the miRNA and the siRNA pathways [14].

Although the identification of conserved regions or motifs between species could indicate functional importance, rapidly evolving genes could escape this trend. We next used FEL (Fixed Effects Likelihood) [39] to explore signals of positive/negative selection at the level of specific sites. This approach estimates the ω rates for each site of a given coding alignment and phylogeny using maximum likelihood methods. We applied this methodology to *loqs* and *loqs2* separately in both *Ae. aegypti* and *Ae. albopictus* (**Figure 3F** and **Supplementary Figure 2B**). We focused on sites that were identified in both species that presumably reflect general trends for each protein. Regarding *loqs*, we observed 35 sites under negative selection that were common between *Ae. aegypti* and *Ae. albopictus*. These results confirm that *loqs* is evolving under strong purifying selection to conserve its functional properties in both species. In contrast, in the case of *loqs2*, we observed only 1 site under negative and 5 sites under positive selection that were common between *Ae. aegypti* and *Ae. albopictus*, 4 of which were located outside of dsRBDs and were mainly located at the C-terminal region (**Figure 3F** and **Supplementary Figure 2B**).

These findings suggest that the C-terminal region of *loqs2* might have evolved under diversifying selection probably towards new protein-protein interactions and functions, while the dsRBDs mostly remained under pressure to maintain dsRNA binding ability. Similarly, comparison between *Drosophila* species show that positive selection is mostly observed outside conserved domains of *Dcr-2, AGO2* and *r2d2* [36]. This intra-protein evolutionary trend could provide novel and yet unknown functions to Loqs2 independent of the constrains of RNAi activity, such as enabling the communication between pathways upon the recognition of specific dsRNAs.

### *loqs2* has unique pattern of expression that is distinct from *loqs* and *r2d2*

Next, we used publicly available RNA-seq data to analyze expression patterns of *loqs, loqs2*, and *r2d2* in different tissues of *Ae. aegypti* and *Ae. albopictus* compared to *loqs* and *r2d2* in *An. gambiae*. We observed that the expression of *loqs2* in both *Ae. aegypti* and *Ae. albopictus* was mostly restricted to reproductive tissues (**Figure 4A**). In contrast, *loqs* and *r2d2* showed ubiquitous expression in all three mosquito species (**Figure 4A**). In addition to reproductive tissues, we also noted high expression of *loqs2* in early embryos (0-4h) followed by a sudden decrease in later stages (24-72h) suggesting that *loqs2* mRNA is maternally deposited in the embryo. In addition, we observed no expression of *loqs2* in larval or pupal stages, suggesting tight regulation of its expression during mosquito development. Overall, the unique expression pattern of *loqs2* gives further support for its neofunctionalization after duplication from an ancestral *loqs* gene.

**Figure 4.**
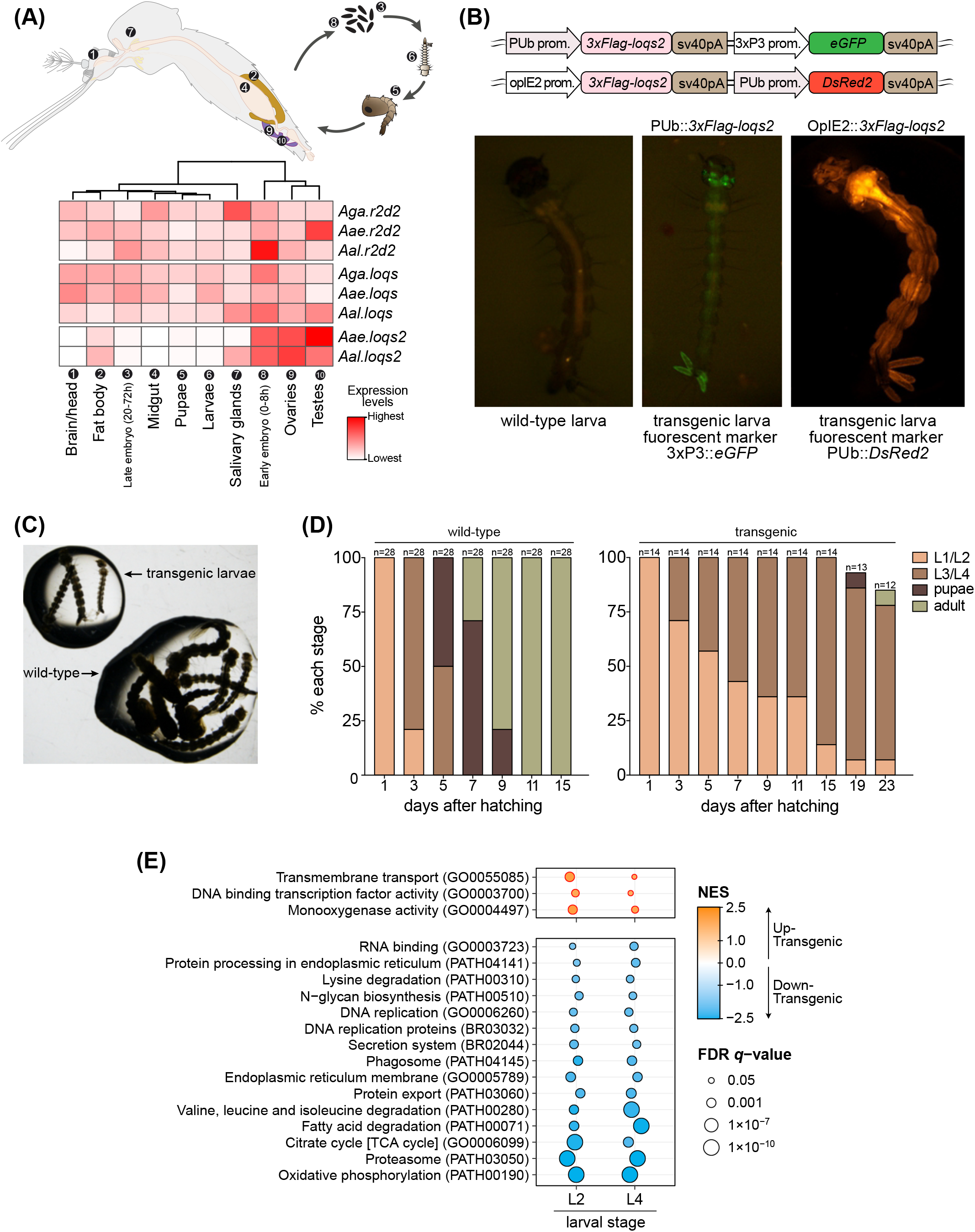
Ectopic expression of *loqs2* in larval stages results in larval growth arrest. **(A)** Heatmap showing the tissue-specific expression of *loqs, loqs2* and *r2d2* among *Ae. aegypti, Ae. albopictus* and *An. gambiae*. Gene expression between tissues was used for hierarchical clustering. White boxes are indicative of no detected mRNA expression. **(B)** Cassettes for ectopic expression of loqs2 under the control of the *Ae. aegypti poly-ubiquitin* (*PUb*) promoter or the baculovirus promoter OpIE2. Fluorescent markers under control of the promoters 3x*P3* or *PUb* were used to drive eGFP or DsRed2 expression as transgenesis markers as indicated. Images are representative of larvae from each transgenic line. **(C)** Wild-type (non-transgenic) sibling larvae reared at the same water container show normal development until pupal stage while transgenic larvae were halted in L2-L3 stages of development at day 5 post-hatching. **(D)** Stacked bar plots comparing development of wild-type and transgenic mosquitoes ectopically expressing *loqs2* under control of the baculovirus promoter OpIE2. **(D)** Gene set enrichment analysis (GSEA) of larvae ectopically expressing *loqs2* compared to wild-type siblings showed a major transcriptional shutdown on the transcription of metabolic pathways in transgenic individuals.

In order to investigate potential functions of *loqs2*, we evaluated if perturbations in *loqs2* expression could impact development in *Ae. aegypti*. For this, we generated two independent transgenic mosquito lines expressing *loqs2* under the control of either the *poly-ubiquitin* (PUb) gene promoter or the baculovirus promoter OpIE2 (**Figure 4B** and **Supplementary Figure 3A**). These promoters drive ubiquitous gene expression at late larval stages of *Aedes* and, in both cases, caused delayed larval development compared to non-transgenic siblings. Most transgenic individuals showed retarded molting with consequent prolonged L3 and L4 larval stages (**Figures 4C,D**). As a comparison, transgenic mosquitoes expressing eGFP or DsRed2 under the control of the same promoters do not show any developmental phenotypes (data not shown). Transcriptomic analysis of transgenic larvae and wild-type siblings at L2 and L4 stages showed significant differences in gene expression between both conditions. In transgenic mosquitoes, gene-set enrichment analysis (GSEA) showed a significant decrease in expression of genes related to metabolism and metabolic pathways, and upregulation of biological processes related to the control of reactive oxygen species (**Figure 4E** and **Supplementary Figure 3B**). Our results highlight the importance of tissue specific control of *loqs2* expression. It is tempting to speculate that the presence of Loqs2 could impact viral tropism since it is absent from tissues well known to host strong arbovirus replication such as the midgut. Loss-of-function experiments will be necessary to validate this hypothesis and better understand tissue-specific roles of Loqs2 in *Aedes* mosquitoes.

## Discussion

Our group has recently reported that the ectopic expression of *loqs2* in the midgut of *Ae. aegypti* mosquitoes, where Loqs2 is naturally absent, enables the control of DENV or ZIKV infection by the mosquito [19]. These results suggest that Loqs2 could have evolved to be a strong regulator of the siRNA pathway, potentially balancing the response triggered by viruses or exogenous and endogenous dsRNAs. Recognition of viral dsRNA is indeed key to the siRNA pathway, and gene duplications of dsRBPs involved in RNAi are commonly observed in insects (**Supplementary Figure 1A**) [40,41]. In the fly model *D. melanogaster, blanks* is a recent paralog exclusive to this species that contains two dsRBDs. *Blanks* is highly expressed in *Drosophila* testes, and knock-out male flies are infertile due to spermiogenesis issues [40,41]. In *D. melanogaster* S2 cells, ablation of Blanks leads to defect in the siRNA pathway, suggesting that Blanks is a co-factor of RNAi. Strikingly, *Dicer-2* knockout flies are fertile but the role of Blanks in fertility seems independent of dsRNA binding, showing the versatility of these proteins [42]. Tracing a parallel with our current work, we observed that *loqs2* is also highly expressed in reproductive tissues of the mosquito. Interestingly, in addition to dsRBPs, duplication of *AGO2* that are only expressed in testis has also been observed in *Drosophila* species [43]. These observations suggest that the siRNA pathway is highly active in the gonads and may affect reproduction in addition to the antiviral defense. Notably, genes affecting host-pathogen interactions and reproduction are both expected to have a strong signal of positive selection.

Of note, Loqs2 presents the same domain organization as Loqs-PD, a *D. melanogaster* Loqs isoform that evolved to be a regulator of the siRNA pathway in this species. As mentioned before, *Ae. aegypti* Loqs has 3 predicted isoforms (Loqs-PA, -PB and -PC) [14]. Interestingly, *Ae. aegypti* Loqs-PC is a predicted isoform that lacks the C-terminal Staufen C domain, similar to *D. melanogaster* Loqs-PD. Since *Ae. aegypti* Loqs-PC has no detectable expression in tissues of the insect, the role of this isoform could have been somewhat replaced by the appearance of *loqs2*. This could have led to the specialization of Loqs by negative selection and the diversification of Loqs2 towards new tissue specific functions. Identification of isoforms lacking the Staufen C domain in other species belonging to *Aedes, Anopheles* and *Culex* genus could provide supporting evidence to this hypothesis. However, except for *Ae. aegypti*, the current genome annotation of these species available in Vectorbase (release 52) only exhibit one transcript correspondent to the long isoform of *loqs* that codes for orthologs of *Ae. aegypti* Loqs-PA. The unavailability of different isoforms is possibly a result of poor annotation of RNAi genes in these species, and further identification of isoforms would require more studies.

An outstanding question is whether the molecular functions of Loqs2 are linked to that of Loqs, especially considering that our data indicate its origin from two independent duplications of the first dsRBD of *loqs*. In *D. melanogaster*, both dsRBDs of Loqs seem to be equally capable of binding dsRNA [25]. Nevertheless, the first dsRBD of *D. melanogaster* Loqs-PD is essential for the cleavage of suboptimal dsRNA substrates by Dicer-2 *in vitro* [44]. The flexibility and length of the amino acid linker connecting both dsRBDs of *D. melanogaster* Loqs seems to allow independent binding of each domain to a dsRNA substrate, with consequent and mobile reversible interaction [25]. In *Aedes* mosquitoes, Loqs has a linker length between dsRBDs D1 and D2 of 29 amino acids while this distance in Loqs2 is only of 15 amino acids. We speculate that the short linker between the two dsRBDs of Loqs2 result in lower flexibility and restrict binding to the same dsRNA molecule. This may have caused a shift in binding affinities to dsRNA substrates of different origin and conferred evolutionary advantages that helped select Loqs2. This hypothesis is also corroborated by the signs of positive selection observed on the linker of Loqs2 (**Figure 3F**). Taken together, we hypothesize that Loqs2 might have evolved to deal with unique dsRNA substrates possibly in response to the evolutionary arms race with viruses.

In the same perspective, we wondered if Loqs2 could bear similar protein-protein interaction features as others RNAi co-factors such as Loqs and R2D2. In *D. melanogaster*, the interaction of different isoforms of Loqs with other partner proteins occur independently of the two dsRBDs. In *D. melanogaster*, Loqs-PA interacts with Dicer-1 exclusively through its Staufen C domain, while Loqs-PD interacts with Dcr-2 through a 37 aa C-terminal region [37,44,45]. These regions are not present in Loqs2 but we observed strong signals of positive selection in its C-terminal region, which could indicate this may be the main site for interaction with protein partners. Notably, binding of both *D. melanogaster* R2D2 to Dcr-2 is also mediated by its C-terminal region and not dependent on the dsRBDs [44,46]. Nevertheless, we cannot rule out that the interaction between Loqs2 and partner proteins such as R2D2 and Loqs in *Ae. aegypti* are indirect and dependent on binding to dsRNA [19].

This work has begun to shed light into the origin and evolution of *loqs2*, an antiviral gene in *Aedes* mosquitoes. More studies are needed to further characterize *loqs2* functions and how its appearance may have shaped the evolution of vector competence in mosquitoes. With the recent advances in genetic-based strategies, *loqs2* seems like a promising target for genetic interventions since regulation of this *Aedes* specific gene could render mosquitoes resistant to virus infection. Strategies such as population replacement using gene-drive to modify mosquito populations to express *loqs2* in the midgut could be an alternative for targeting the transmission of arboviruses.

## Methods

### *loqs* and *loqs2* sequence retrieval from publicly available high-throughput sequencing data of different species

Public high-throughput RNA-seq and DNA-seq libraries from various mosquitoes (**Supplementary table 1**) were obtained from the NCBI/SRA database and mapped to the *Ae. aegypti* and *Ae. albopictus* reference genomes (Vectorbase release 52) [47] using STAR v2.7.9a [48]. To overcome the expected divergence between the RNA-seq data and the reference genomes, we allowed a maximum alignment mismatch percentage of 50%. Mapping coverage comprising regions of *loqs* and *loqs2* were manually inspected using IGV v2.8.3 [49] to avoid false positives. Data was imported into R v4.1.0 [50] using in-house scripts and plots were generated using the R package ggplot2 v3.3.3.

### Transcriptomic analyses of tissue-specific libraries

To quantify the expression profiles of *loqs, loqs2* and *r2d2* among different tissues from *Ae. aegypti, Ae. albopictus* and *An. gambiae* mosquitoes, public RNA-seq libraries were obtained at NCBI/SRA as described above. Subsequently, reads were mapped to the decoyed reference transcriptome of each species (Vectorbase release 52) using Salmon v1.5.2 [51]. Quasi-mapping quantifications were imported into R v4.1.0 [50] and data normalization was performed using the packages EdgeR v3.34.0 and TMM [52,53]. Plots were generated using the R package ggplot2 v3.3.3.

### Phylogenetic analyses

For phylogenetic analyses we used a set of mosquitoes and flies to infer the evolutionary history of *loqs2* at three different evolutionary ranges: *(i)* the dsRBP orthologs (InterPRO accession IPR014720) [54], *(ii)* the close related evolutionary history of *loqs2*, including *loqs* and *r2d2* and, *(iii)* the *loqs2, loqs* and *r2d2* single domain evolutionary history. Input sequences were aligned using either PRANK_+F_ (strategies *i* and *iii)* [55] or MAFFT-E-INS v7 [56] (strategy *ii)* and the substitution model and phylogenetic relationships were inferred using IQ-tree2 [57]. For all cases, the maximum-likelihood consensus tree was generated with 1000 ultra-fast bootstraps iterations. All trees were rooted at the midpoint using Figtree v1.4.4 [58] for visualization purposes. For the dsRBP evolutionary range, we used VectorBase (release 52) to retrieve amino acid sequences of the dsRBPs (InterPRO accession IPR014720), including Dicer-2. The WAG amino acid exchange matrix under FreeRate heterogeneity model with 5 categories (WAG+R5) was determined to be the best fit model based on the Bayesian information criterion (BIC). For the *loqs2* closely related tree we used the coding sequences from the orthologs belonging to the same set of species of strategy *i*. The KOSI07 empirical codon model with discrete Gamma model with 4 rate categories (KOSI07 + G4) was determined to be the best fit model based on the BIC. For the single domain tree, we used the coding sequences of each domain from *loqs2, loqs* and *r2d2* belonging to *Ae. aegypti* and *Ae. albopictus*. The KOSI07 empirical codon model with amino-acid frequencies given by the protein matrix (KOSI07 + FU) was determined to be the best fit model based on the BIC.

### Loqs and Loqs2 dsRBDs tridimensional modeling and alignment, amino acid sequence comparison

Homology models of the dsRBDs of *Ae. aegypti* and *Ae. albopictus* Loqs and Loqs2 were built using a template-based method. The *D. melanogaster* Loqs-PD dsRBD1 structure (PDB id: 5NPG) was used as the structural template to model the dsRBD1 of Loqs and the dsRBDs of Loqs2. For Loqs dsRBD2 models, the *D. melanogaster* Loqs-PD dsRBD2 structure (PDB id: 5NPA) was used as template [25]. Models were made with MODELLER v9.24 [59] using the automodel class. In each case, 100 models were produced, and the model with lowest MODELLER DOPE score was selected. Quality of each selected model was checked by the analysis of its Ramachandran plot built with PROCHECK [60]. All selected models presented at least 88% of their residues located in the most favored regions of the plot and no more than 1 residue located in a disallowed region. Model visualization was done using pymol v2.4.0 [61]. For structural comparisons, the modeled *Ae. aegypti* and *Ae. albopictus* Loqs and Loqs2 dsRBDs were compared to each other using the DALI server [26]. For the amino acid sequence alignment and comparison, the sequences of *Ae. aegypti* and *Ae. albopictus* Loqs and Loqs2 dsRBDs and Staufen C amino acid sequences were pairwise aligned with BLASTp under the BLOSUM62 substitution matrix [62]. Score and coverage of the pairwise alignments were summarized in a correlation matrix plotted using R v4.1.0 and the package ggplot2 v3.3.3.

### Evolutionary analyses using a phylogenetic framework

We used the phylogenetic tree including *loqs, loqs2* and *r2d2* as evolutionary hypothesis to perform different tests of selection based on the rates of nonsynonymous and synonymous substitutions (*ω)*. A test for relaxation or intensification of selection was performed with RELAX [28] implemented in the datamonkey server [63]. RELAX uses the parameter *k* to test for relaxed (*k* < 1) or intensified (*k* > 1) selection. RELAX tests for statistical significance (*p-value* < 0.05) by comparing a null model (*k* is fixed to 1) to an alternative model (*k* is a variable parameter) using a likelihood ratio test (LRT). Paired comparisons were made between the *Aedes* orthologs of *loqs* and *loqs2* as foreground branches and *loqs, loqs2* and *r2d2* as background branches. All comparisons were made including and excluding the internal branches of the tree. Positive selection at the branch level was evaluated using the branch-site-model implemented in CodeML (CodeML-BSM) [29] and automated in EasyCodeML [64], and abSREL [30] implemented in the datamonkey server. Both methods estimate *ω* rates at branch- and site-level of the test branches though under different assumptions. CodeML-BSM uses a model that allows positive selection along the foreground branches and sites only permitting four possible configurations of branch-specific rate parameters [31]. abSREL, infers the optimal number of *ω* classes and models their heterogeneity allowing for all possible configurations of branch-specific rate parameters. Both methods test for selection by using a LRT to compare the adaptive model to a null model that only allows neutral and negative selection. We used both methods to test for selection acting on the branches of *loqs* or *loqs2* before the interspecific diversification of *Aedes, Culex* or *Anopheles* genus. FEL (Fixed Effects Likelihood) [39], implemented in the datamonkey server, was used to evaluate the selective forces acting at the sites-level on each of the *Aedes loqs* and *loqs2* paralogs. This approach estimates the site-specific nonsynonymous (dN) and synonymous (dS) substitution rates using a maximal likelihood approach. Basically, FEL uses the entire dataset to optimize branch lengths and nucleotide substitution parameters, then fits a MG94xREV model to each codon site and infers site-specific dN and dS substitution rates. Finally, it fits a neutral and selection model to every codon and calculates a standard LRT to decide if the site is evolving non-neutrally.

### Evolutionary analyses using population genomics

We used public exome data of an *Ae. aegypti* population from a Senegal forest area [34] (SRA project: SRP092518). Low quality reads were removed and sequencing adaptors were trimmed using trimmomatic v0.39 [65], overlapping reads were merged with FLASH v1.2.11 [66] and both paired- and single-end pseudo-reads were mapped to the *Ae. aegypti* reference genome (Vectorbase release 52) using BWA-mem2 v.2.0pre2 [67]. SAMtools v1.13 [68] was used to merge and sort the paired- and single-end alignments into a single BAM file [68]. Read duplicates were marked with Picard v2.21.5 [69] and variant calling was performed with GATK v4.1.4.1 [70] with a heterozygosity prior of 0.003125 as reported by Redmond et al (2020) for *Ae. aegypti*. SNP calling was performed only for *loqs, loqs2, Dicer1* and *Dicer2* genomic loci including flanks of 2-Mb. Resultant SNP data was hard filtered with VCFtools v0.1.17 and indels were removed [71]. To determine the derived alleles, we performed the same SNP calling pipeline for *Aedes mascarensis* (SRA accession: SRR9959083) whole genome sequence. Finally, we used bcftools and GATK v4.1.4.1 to transform the SNP variants into a fasta sequence and used a custom python script to create a multi-fasta file for *loqs* or *loqs2* coding sequences. These files were used to calculate the nonsynonymous (*Ka)* to synonymous *(Ks)* substitutions using the iMKT web server [33].

### Mosquito transgenesis and developmental assays

Embryo microinjection was performed with small modifications, as previously described [19]. Briefly, freshly laid *Ae. aegypti* eggs were aligned in parallel against the internal side of a U-shaped nitrocellulose membrane in contact with an overlaying filter paper soaked in demineralized water. Aligned eggs rested in a humid chamber for 30–60 min after alignment, until the eggs turned dark grey. Mixes containing 100 ng/μl of a *piggyBac* transposase helper-plasmid and 400 ng/μl of either plasmid carrying *piggyBac*-flanked cassettes (**Figure 4B**) were diluted in 0.5 × PBS and injected at the embryo posterior pole under a Nikon Eclipse TE2000-S inverted microscope, using a Femtojet injector (Eppendorf) and a TransferMan NK2 micromanipulator. Before allowed to dry, microinjected eggs were kept for 2 days diagonally in a container with 1-cm-deep water. The filter paper had direct contact with water, which kept the eggs moist by capillarity [72]. Surviving eggs were hatched under vacuum and larvae showing constitutive expression of fluorescent markers were kept for further crossing with wild-type individuals.

### Poly-A selection, high-throughput RNA-seq library construction, and transcriptomic analysis

Total RNA from 5 to 10 individual larvae were extracted using TRIzol (Invitrogen) following the recommended protocol. RNA integrity was verified using the 2100 Bioanalyzer system (Agilent). mRNA libraries were constructed using the kits NEBNext Poly(A) mRNA Magnetic Isolation Module and NEBNext UltraTM II Directional RNA Library Prep Kit for Illumina (New England BioLabs) following the manufacturer protocol. Indexed libraries were pooled and sequenced at the GenomEast sequencing platform at the Institut de Génétique et de Biologie Moléculaire et Cellulaire of Strasbourg, France. Sequenced reads with an average quality score above phred 25 had adaptors removed using Trimmomatic v0.39 and were further mapped to the decoyed transcriptome of *Ae. aegypti* (Vectorbase release 52) using Salmon v1.5.2 [51,65]. Quasi-mapping quantifications were imported into R v4.1.0 (R Core Team 2021) and data normalization was performed using the packages EdgeR v3.28.1 and TMM [52,53]. Ranked lists of gene expression for each comparison (transgenic against wild type larvae) was used as input for Gene Set Enrichment Analysis (GSEA) [73] using the R package fgsea v1.12.0 [74] and in-house developed gene-sets comprising Gene Ontology annotation, KEGG pathways, and genes of interest. Sets with adjusted FDR *q*-value < 0.1 were considered in our analysis and redundancy was removed if overlapping genes within given gene-sets were higher than 90%.

## Acknowledgments

We thank all members of the Marques laboratory and the M3i unit - Insect Models of Innate Immunity for suggestions and discussions.

## Author contributions

Conceptualization: CFEC, JTM, RPO

Funding acquisition: JTM

Methodology: CFEC, MFR, FVF, EM, RC, JTM, RPO

Investigation and data curation: CFEC, MFR, AB, FVF, EM, RC, JTM, RPO

Formal analysis and visualization: CFEC, MFR, FVF, RC, JTM, RPO

Writing: CFEC, MFR, JTM, RPO

## Data availability

Transcriptome libraries from this study have been deposited on the Sequence Read Archive (SRA) at NCBI. Other publicly available RNA-seq data sets were obtained from SRA. Accession numbers are provided in **Supplementary Table 1**.

## Supplementary data

**Supplementary figure 1.**
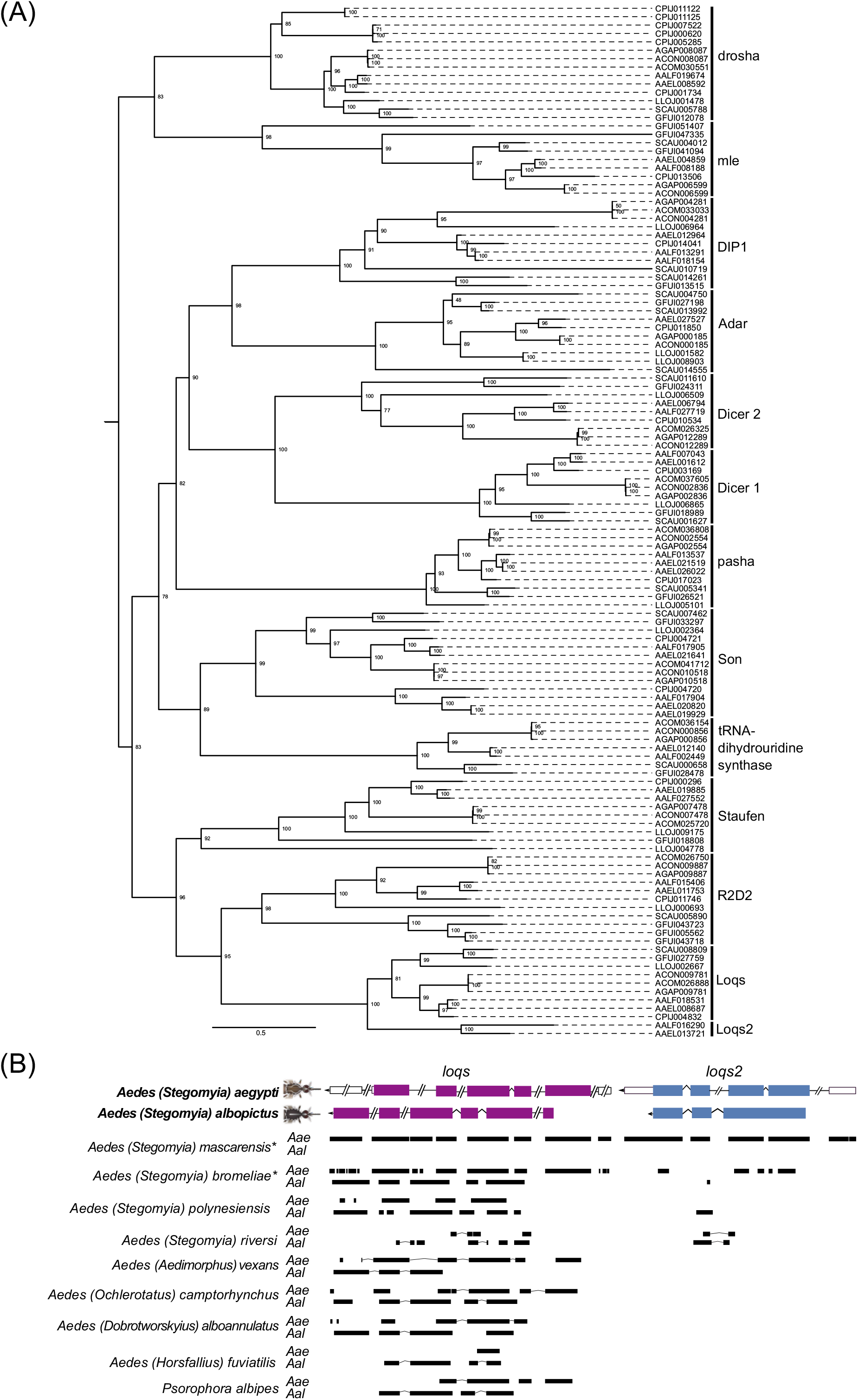
Phylogenetic tree of the dsRNA binding proteins found in the genomes of different mosquitoes and flies, and mapping coverage of RNA-seq and whole-genome-seq data from different *Aedini* species mapped to the *Ae. aegypti* and *Ae. albopictus* reference genomes. **(A)** Terminals correspond to the VectorBase accession numbers. Tree was rooted at the midpoint for visualization purposes. Node values correspond to the result of 1000 ultra-fast bootstrap iterations. **(B)** *loqs* and *loqs2* exons are colored in purple and blue respectively. Black bars represent the read coverage along gene sequence. Lines represent intronic regions of both gene sequences and reads.

**Supplementary figure 2.**
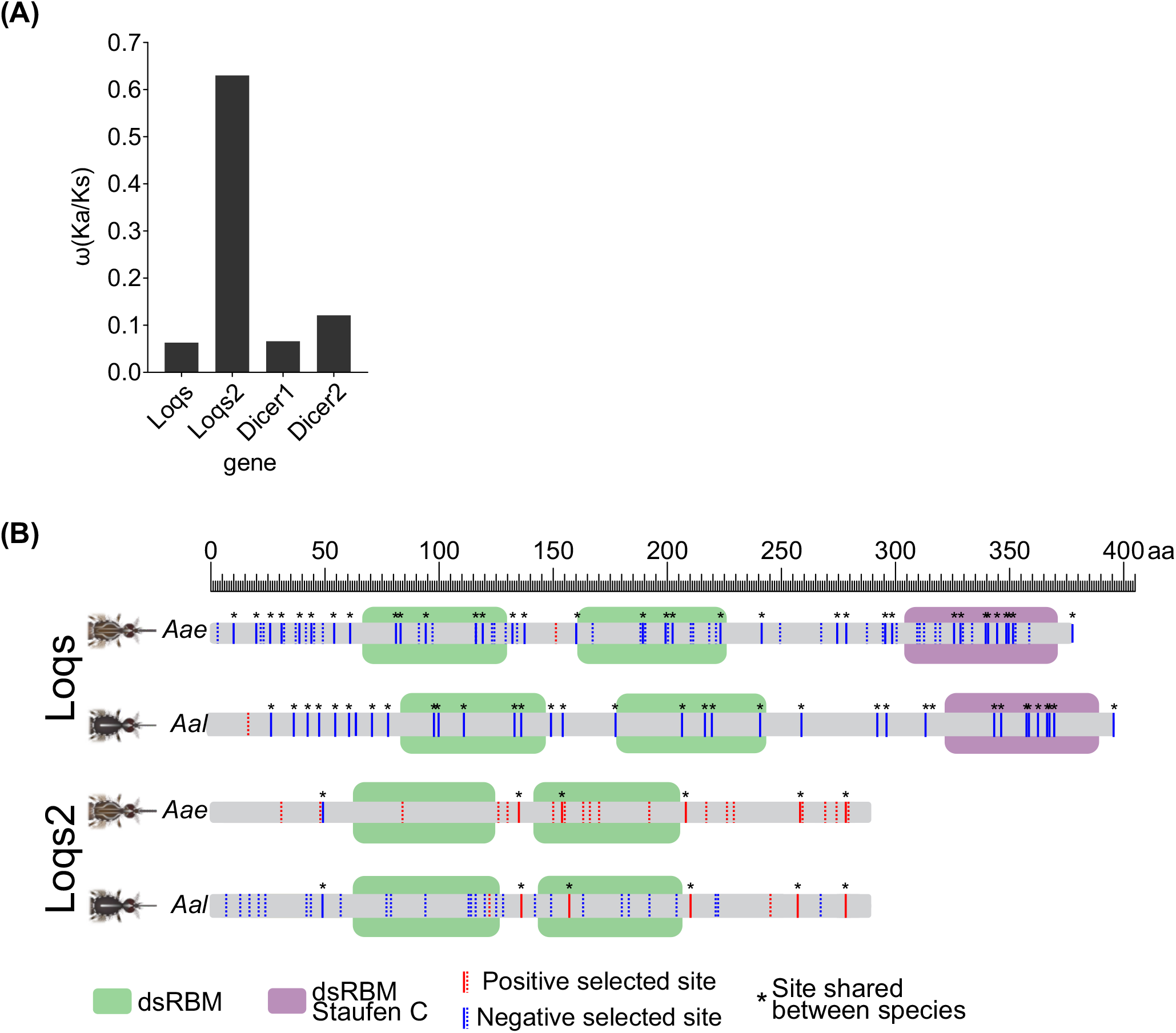
Rate of nonsynonymous divergence (*Ka*) to the proportion of neutral fixations *(Ks)* (ω) among *loqs, loqs2, Dicer-1* and *Dicer-2* genes, and complete comparative scheme between *Ae. aegypti* and *Ae. albopictus* Loqs and Loqs2 proteins. **(A)** Analysis was performed comparing a Senegal Forest *Ae. aegypti* population and *Ae. mascarensis*. **(B)** Solid lines indicate sites shared between species, as shown in **Figure 4B**, while specific sites are indicated with dotted lines.

**Supplementary figure 3.**
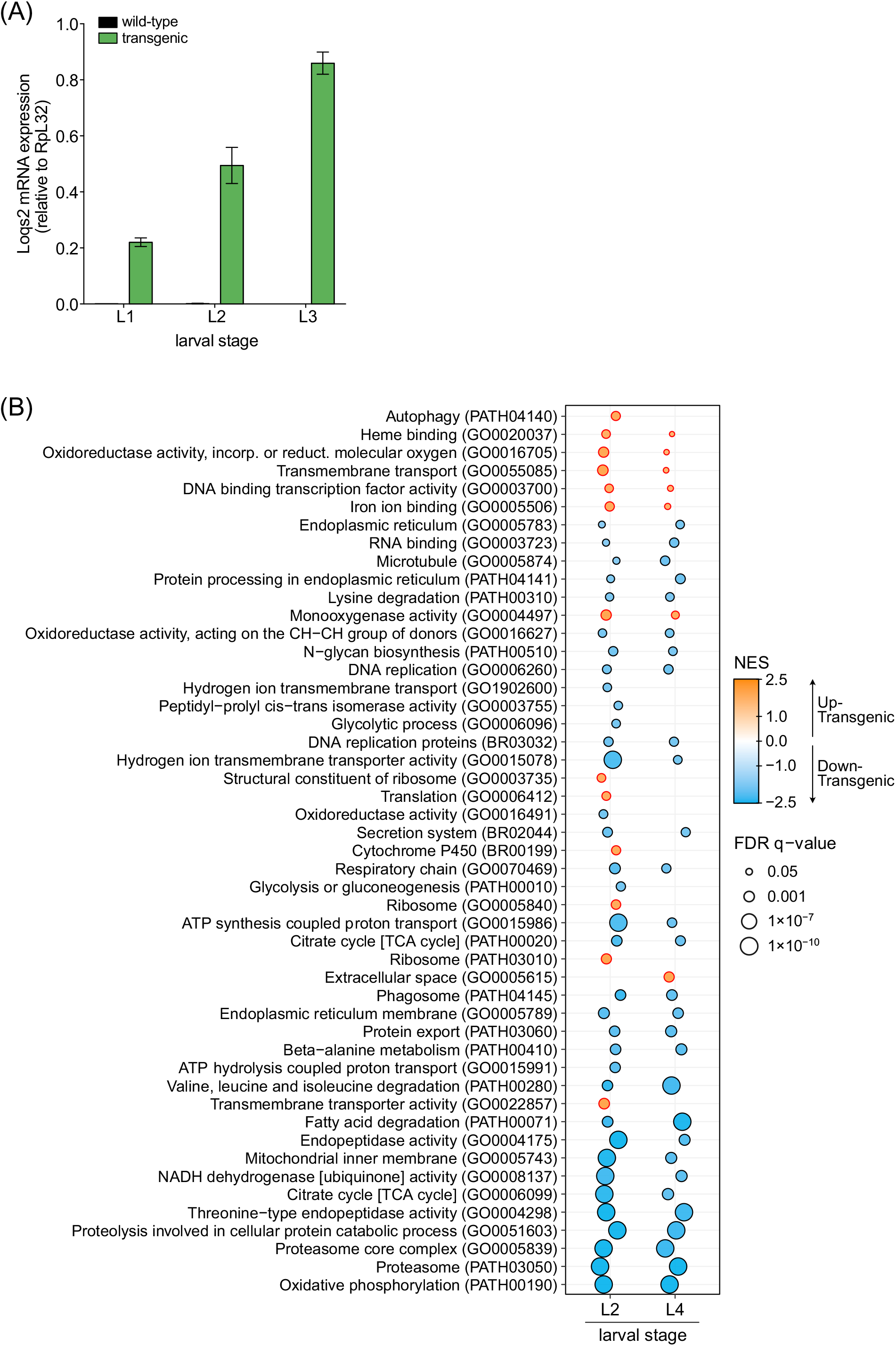
Ectopic expression of *loqs2* in larval stages lead to major metabolic shutdown. **(A)** Expression levels of *loqs2* in transgenic or wild-type sibling larvae quantified by RT-qPCR. **(B)** GSEA analysis plot showing extended gene sets prior redundancy removal relative to **Figure 4E**.

**Supplementary table 1. SRA identifiers of the libraries used in this study**. Excel spreadsheet file.

